# STING agonist-mediated endothelial cell activation drives NK cells and neutrophils-dependent pulmonary inflammation

**DOI:** 10.64898/2026.03.10.710764

**Authors:** Chen Chen, Yaqi Zhao, Fengyu Du, Rongchen Liu, Xinhang Zheng, Shanna Wu, Yiting Wang, Fengru Qiu, Liying Chen, Runqiu Chen, Fanglin Li, Likun Gong, Yiru Long

**Author notes:** **Correspondence:** Likun Gong or Yiru Long.

## Abstract

Stimulator of interferon genes (STING) agonists and derivative molecules have been extensively developed for tumor immunotherapy. However, systemic exposure toxicity risks have constrained clinical trial progression and even threatened patient lives. Currently, systematic toxicity assessments for STING agonists remain lacking, with the mode of action for major organ injury yet to be elucidated. Here, we focused on STING agonist-induced lung injury, revealing that systemic administration of STING agonists caused pulmonary hemorrhage, inflammatory alterations, and respiratory dysfunction. Through single-cell RNA sequencing and immune deletion studies, we found that lung endothelial cells could be stimulated by STING agonists and then secreted chemokines and IL-15 to recruit and activate NK cells. NK cells could induce endothelial cell apoptosis via IFN-γ. Tbx21^+^ NK subpopulations, which activated by endothelial cells, could produce chemokines to recruit neutrophils. Neutrophils secreted IL-1β through positive feedback pathways and form neutrophil extracellular traps during lung injury. This study elucidates the critical role of the endothelial cell-NK cell-neutrophil axis in mediating STING agonist-associated pneumonia, offering insights for developing intervention strategies for STING agonist toxicity.

## Introduction

The cyclic GMP-AMP synthase (cGAS)-stimulator of interferon genes (STING) pathway detects pathogen DNA or damage-associated molecular patterns (DAMPs) DNA, serving a vital function in the host’s defense against infections and malignancies. cGAS, as a double-stranded DNA sensor, catalyzes the production of 2’3’ cyclic GMP-AMP (cGAMP)-the endogenous ligand for STING-upon activation [1]. Subsequently, activated STING transfers from the endoplasmic reticulum to the Golgi apparatus and promotes downstream effects such as interferon (IFN) signaling, inflammasome activation, and light-chain 3B (LC3B) lipidation through processes including TBK1-IRF3, NFκB, and proton leakage [2].

Activation of the STING signaling promotes the antitumor effects of dendritic cells (DCs), CD8^+^ T cells, natural killer (NK) cells, etc[3]. Besides, studies have further proposed that STING activation in vascular endothelial cells can remodel the tumor vasculature [4] and alter the structure of the tumor immune microenvironment in “cold” tumors [5].

The anti-tumor effects of STING agonists were first observed in the cGAMP [6]. Studies have demonstrated that injecting cGAMP into the glioma significantly reduces tumor volume and improves survival rates of tumor-bearing mice in a STING-dependent manner [7].

Meanwhile, amidobenzimidazole (ABZI)-based analogs were designed to improve systemic delivery, which can bind to the C-terminal domain of STING and enhance the biding affinity. The representative one is diABZI from linked ABZIs, which showed a potent effect in ameliorating the affinity to STING and inducing the secretion of IFN-β in human peripheral blood mononuclear cells (PBMCs). Administration of diABZI in mice bearing CT26 colorectal tumors resulted in significant tumor inhibition and enhanced survival, with 80% of mice being tumor free [8]. Considering of the superior antitumor effects, numerous STING agonists have now entered clinical development phases. For example, ADU-S100 [9] and E7766 have been employed for treating advanced solid tumors or lymphoma [10]. Beyond this, antibody-drug conjugates (ADCs) targeting STING agonists have also emerged as a focal point in antitumor drug development. Examples include Takeda Pharmaceutical Company’s STING agonist-linked ADC, TAK-500, which targets CCR2 [11].

However, multiple studies have raised concerns regarding the safety of STING agonists.

The injection of small-molecule STING agonists may lead to rapid systemic distribution, thereby posing risks of uncontrolled inflammation and cytokine storms, tissue toxicity, and autoimmune damage [12]. Chronic STING activation may also persistently stimulate cytokine production, thereby fostering an inflammatory tumor microenvironment (TME) that promotes tumor progression [13]. Meanwhile, Mersana’s STING agonist-conjugated antibody-drug conjugate XMT-2056, targeting HER2, has experienced a Grade 5 (fatal) serious adverse event (SAE) in its Phase I clinical trial. The company then has announced a voluntary suspension of the trial [14]. In non-human primates preclinical studies of E7766, participating animals exhibited multiple respiratory adverse reactions including dyspnea, hemoptysis, hypoxia, alveolar hemorrhage, pulmonary embolism, and pulmonary oedema [10]. It is evident that the toxicity issues associated with STING agonists have resulted in a narrow therapeutic window during clinical trials, thereby hindering dose escalation and preventing trials from achieving efficacy endpoints.

And recent studies increasingly indicate that activation of the cGAS-STING pathway promotes or exacerbates the development of pneumonia. For instance, following DNA recognition by macrophages, the STING pathway is activated, triggering IL-6 release which subsequently activates fibroblasts and intensifies airway obstruction [15].

Safety concerns regarding STING agonists, such as pulmonary toxicity, have severely limited their clinical development. Therefore, in this study, we focused on systematically evaluating the pulmonary toxicity risk of STING agonists, identifying the characteristics of lung injury induced by STING agonists, and elucidating the underlying mechanisms.

## Results

### Systemic administration of STING agonists causes lung injury in mice

Analysis of human single-cell databases revealed that STING is highly expressed in lung tissues compared to other organs (Fig S1A). This led us to hypothesize that the lung is the most likely target organ for STING agonist toxicity. Thus, we administered the human-mouse cross-reactive STING-specific agonist diABZI [8] to mice and measured inflammatory cytokine expression in lung, liver, kidney, and intestinal tissues. Lung tissues exhibited the most significantly upregulated transcription levels of tumor necrosis factor-alpha (TNF-α), interleukin-1 beta (IL-1β), and IFN-γ (Fig S1B).

To holistically evaluate the in vivo toxicity responses of diABZI, a concentration gradient dosing regimen was established. Following intraperitoneal diABZI administration, mice exhibited significant weight loss alongside elevated lung coefficients and wet-to-dry weight ratios (Fig S1C, Fig 1A and B), preliminarily confirming STING agonist-induced pulmonary injury. To confirm that diABZI-induced pneumonia is indeed caused by STING activation, we administered natural ligand of STING, 2’,3’-cGAMP to mice in vivo. The results mirrored those observed with diABZI administration: the mice exhibited weight loss and elevated expression of inflammatory cytokines in lung tissue (Fig S1D). Based on the the above indicators, a 2 mg/kg dose was selected for subsequent in vivo assays. Further assessment of pulmonary function during treatment revealed reduced peak expiratory flow rates and increased respiratory rates (Fig 1B), indicating ventilatory dysfunction and respiratory impairment. Histopathological examination revealed hemorrhage, congestion, disrupted alveolar architecture, and increased immune cell infiltration in the lungs of diABZI-treated mice (Fig 1C). Surfactant protein A (SP-A), which typically regulates alveolar gas exchange, exhibits serum level elevation associated with pulmonary tissue injury [16, 17]. Western blot analysis demonstrated a significant increase in serum SP-A levels 48 hours post diABZI administration (Fig 1D).

**Fig. 1.**
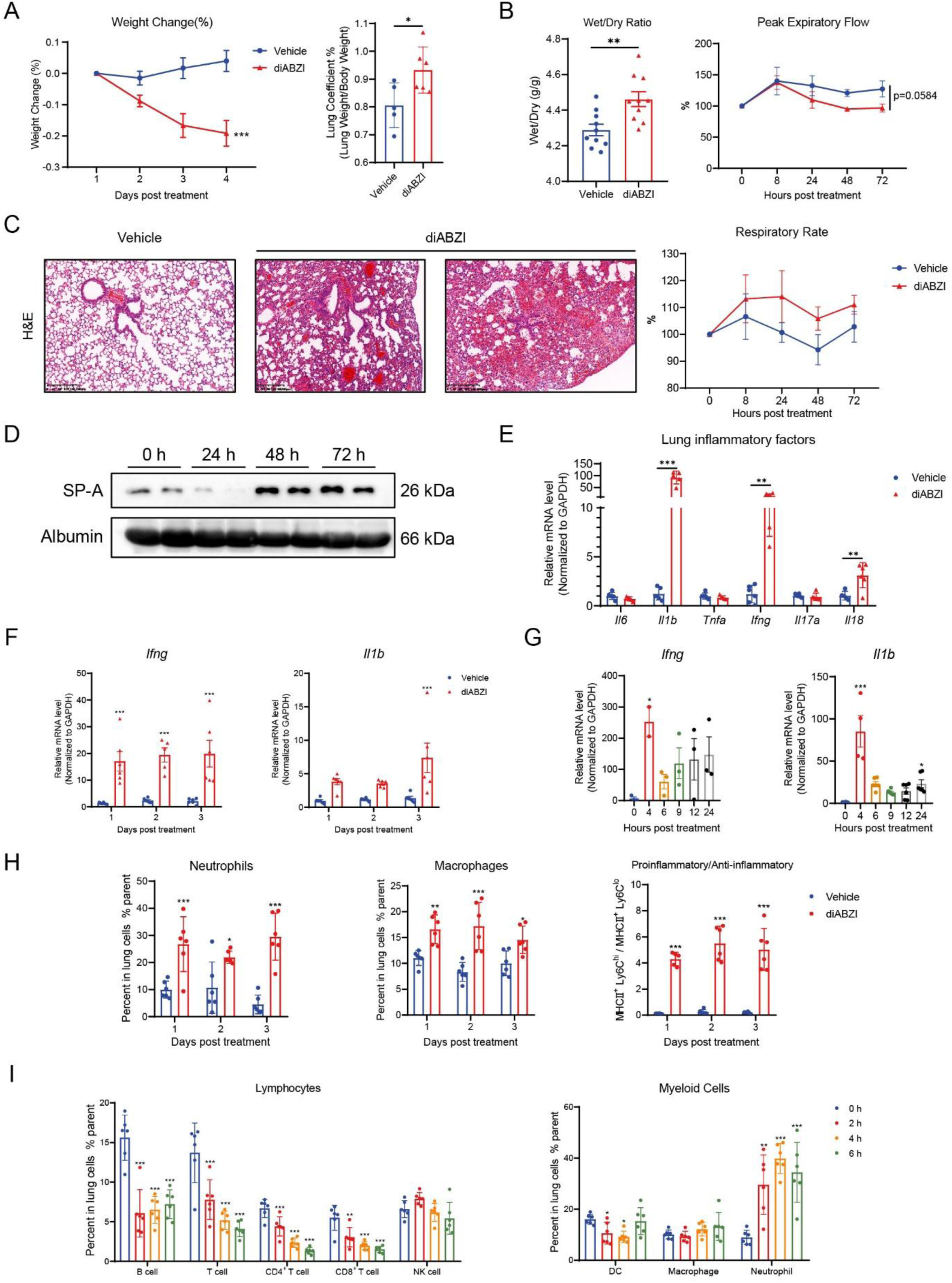
Systemic administration of STING agonists causes lung injury in mice. Female C57BL/6 mice received intraperitoneal injections of 2 mg/kg diABZI every day for three days. (**A**) Body weight change percent of mice was monitored after diABZI treatment from day1 to day4 and lung coefficient was detected on day4 (n=6). (**B**) Lung wet/dry weight ratio was detected on day4 and mice peak expiratory flow and respiratory rate was monitored during day1 to day4 (n=6). (**C**) Representative H&E staining images of lung tissues from mice. Scale bar, 200 μm. (**D**) Western blot analysis of surfactant protein A proteins in mouse serum. (**E-G**) Transcription levels of multiple proinflammatory factor in mouse lung tissues were detected (n=6). (**H**) Flow cytometric analysis of neutrophil macrophage M1-like and M2-like macrophages in the lung tissues (n=6). (**H**) Flow cytometric analysis of lymphocytes and myeloid cells in the lung tissues (n=6). The data are presented as the mean ± SEM. * p < 0.05; ** p < 0.01; *** p < 0.001; ns not significant by ns not significant by unpaired t test or ANOVA followed by Tukey’s multiple comparisons test.

Concurrently, the transcriptional levels of inflammatory cytokines IL-1β, IFN-γ, and IL-18 were substantially upregulated, rapidly peaking within 4 hours post-administration.

Following a decline at 6 hours, levels gradually rebounded to reach a plateau at 24 hours, maintaining elevated expression thereafter (Fig 1E-G). Similarly, in vivo administration of cGAMP also resulted in a significant increase in IFN-γ and IL-1β in lung tissues (Fig S2). Flow cytometry analysis revealed a rapid and significant reduction in the proportion of T/B lymphocytes within the pulmonary immune microenvironment following diABZI administration. Regarding myeloid cells, the STING agonist induced increased neutrophil and macrophage infiltration, reaching a plateau within 24 hours, concurrent with enhanced pro-inflammatory polarization of pulmonary macrophages (Fig 1H and I).

In summary, STING agonists possessed significant pulmonary toxicity risks, potentially leading to pathological lung injury, immune dysregulation, and impaired respiratory function.

### Single-cell sequencing reveals alterations and interactions in pulmonary cell populations post STING activation

To elucidate the alterations in cell population proportions, gene expression changes, and cellular interactions induced by STING agonists, single-cell RNA sequencing (scRNA-seq) was performed on mouse lung tissue collected at 0, 4, 6, 12, and 24 hours post intraperitoneal administration of diABZI. Following t-SNE dimensionality reduction, the data were identified as 12 distinct cell populations. T cells and B cells exhibited markedly reduced proportions following administration, whereas neutrophil and macrophage proportions significantly increased (Fig 2A), consistent with flow cytometry results (Fig 1H and I).

**Fig. 2.**
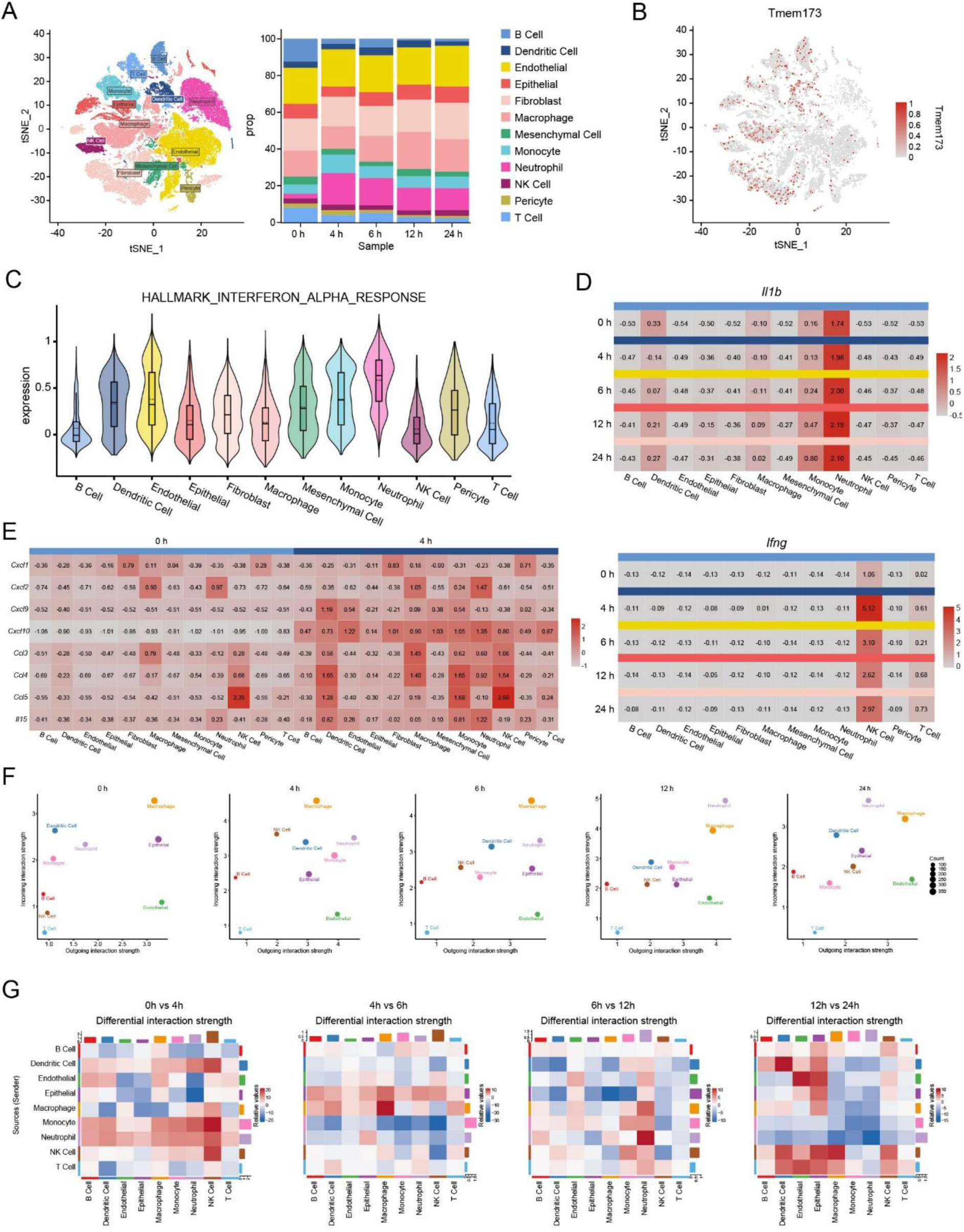
Single-cell sequencing reveals alterations and interactions in pulmonary cell populations during inflammation. (**A**) t-SNE plots and relative proportion of the indicated cell types for scRNA-seq data of lung samples. (**B**) *Tmem173* expression in t-SNE plots. (**C**) Violin plot of gene expression scores for the HALLMARK INTERFERON ALPHA RESPONSE pathway across cell populations. (**D**) Heatmap of IL-1β and IFN-γ expression levels in different cell populations at various time points (**E**) Heatmap of chemokine expression levels in various cell populations at 0 and 4 hours. (**F**) Scatter plots visualized the primary senders and receivers of cellular communication. The x-axis and y-axis respectively represent the total outgoing or incoming communication probabilities associated with each cell population. The size of each point indicates the number of relationships (both outgoing and incoming) with each cell population. (**G**) Heatmap of relative signal strength at different time points (red indicates an increase compared to the previous time point, blue indicates a decrease).

Notably, NK cells, macrophages, T cells, and B cells exhibited high STING expression at rest (Fig 2B). Following STING agonist administration, the IFN-α response pathway was substantially activated in endothelial cells, neutrophils, and monocytes, with increased gene expression (Fig 2C). Previously, we demonstrated that diABZI induced elevated expression of pulmonary inflammatory cytokines IL-1β and IFN-γ (Fig 1E). Further analysis of scRNA-seq data revealed that myeloid cells, particularly neutrophils, constitute the primary IL-1β-secreting population, peaking at 12 hours post treatment. Conversely, NK cells predominantly secreted IFN-γ, peaking at 4 hours; in contrast, other cell populations exhibited negligible IFN-γ secretion (Fig 2D). Combined with scRNA-seq data, flow cytometry results indicate significant alterations in the proportion of pulmonary immune cell populations within 4 hours post-administration.

To investigate the mechanisms underlying changes in the proportion of pulmonary immune cell populations, we analyzed levels of CC and CXC subfamily chemokines. Within 4 hours post-administration, *Cxcl1*, *Cxcl2*, *Cxcl9*, *Cxcl10*, *Ccl3*, *Ccl4*, and *Ccl5* were significantly upregulated (Fig. S3A). Analysis of these chemokine expression levels at 0 and 4 hours by cell population revealed that multiple cell types upregulate chemokine expression following administration. Specifically, DCs, endothelial cells, NK cells, neutrophils, and monocyte-macrophages exhibited markedly elevated chemokine expression (Fig. 2E). This suggests potential interactions and chemotaxis between these cell types following STING agonist administration.

To validate this hypothesis, we employed CellChat for scRNA-seq data analysis and visualization of cellular communications. Results demonstrated that both the number and strength of cellular communication rapidly increased following drug administration, peaking at 4 hours. Subsequently, communications gradually diminished at 4 hours yet remained elevated compared to resting conditions (Fig. S3B). Scatter plots visualizing signal input and output probabilities across cell populations at different time points revealed that NK cell communication probability substantially increased within 4 hours, while neutrophil and endothelial cell output probabilities significantly rose. At 12 hours, neutrophil signal exchange probability peaked, with intercellular communication probabilities subsequently declining post-12 hours (Fig. 2F). Further analysis of specific interacting cell populations and communication intensity revealed that between 0-4 hours, neutrophils and endothelial cells transmitted signals to NK cells, with NK cells exhibiting the strongest signal reception intensity; between 4 and 6 hours, the strength of signals transmitted by epithelial cells increased, primarily received by NK cells, neutrophils, and macrophages; at 12 hours, neutrophils replaced NK cells as the cell population receiving the strongest signals, primarily from NK cells, macrophages, and self-activation; compared to 12 hours, at 24 hours, signal exchange from non-immune cells, including endothelial and epithelial cells, was enhanced (Fig. 2G).

Taken together, we hypothesized that after STING agonist administration, endothelial cells may recruit and activate NK cells in the early phase, and then NK cells may secrete IFN-γ and other chemokines, thereby acting upon neutrophils to enhance their infiltration and IL-1β secretion.

### Endothelial cells can be directly activated by SITNG agonists

Following systemic administration, pulmonary vascular endothelial cells constitute the primary line of direct contact with STING agonists. Analysis of differentially expressed genes and IFNα response pathway genes in pulmonary endothelial cells revealed substantial pathway activation within 4 hours, concurrent with upregulation of chemokines *Cxcl9*, *Cxcl10*, *Ccl4*, *Ccl5*, and classical NK cell activation factors *Il15* (Fig. 3A-C).

**Fig. 3.**
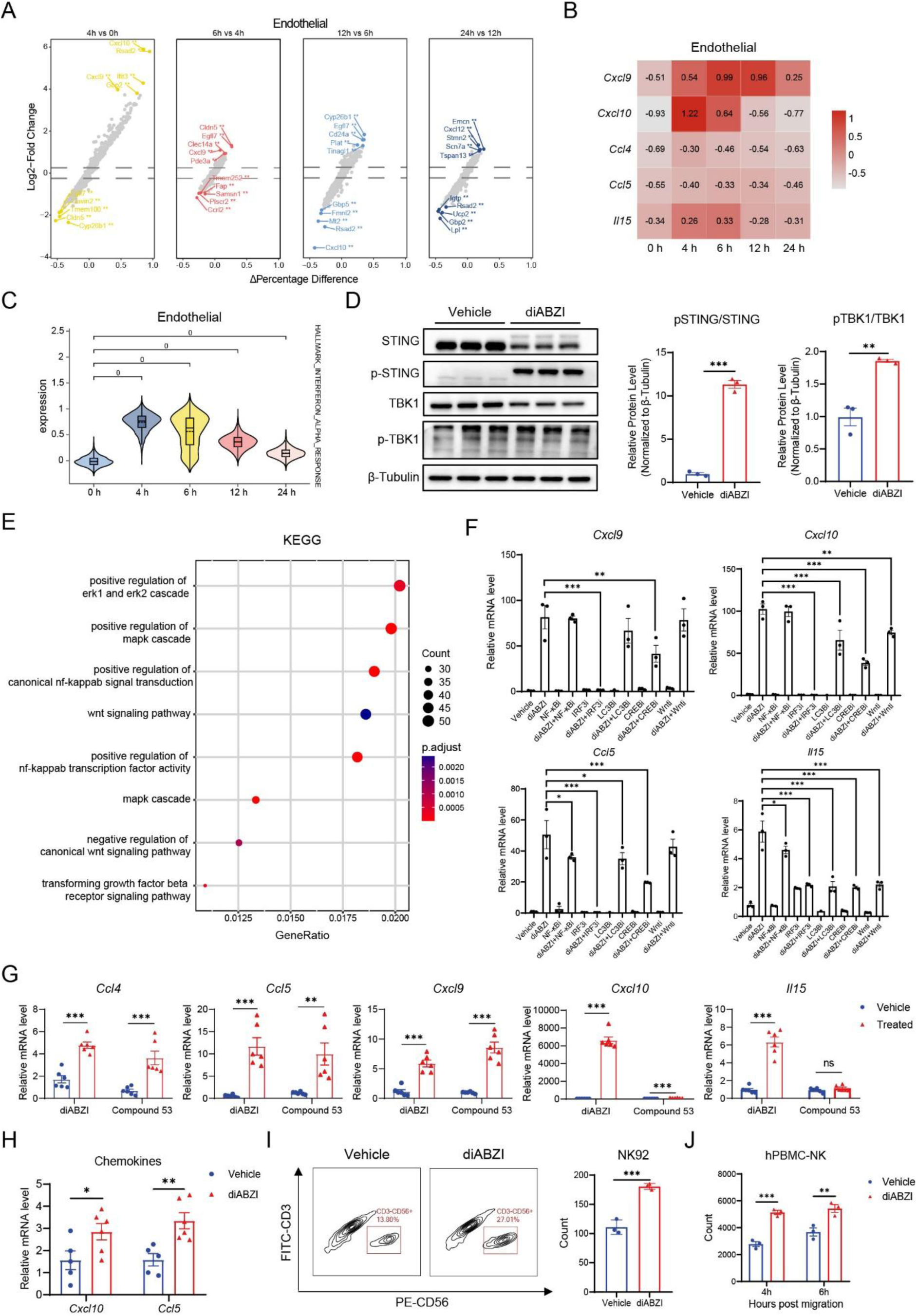
Endothelial stimulated by STING agonist secret chemokines and cytokine to recruit and activate NK cells. (**A**) Volcano plot of gene expression differences of endothelial cells at different time points from scRNA-seq data of lung samples. (**B**) Heatmap of chemokines expression level of endothelial cells. (**C**) Violin plot of gene expression scores for the HALLMARK INTERFERON ALPHA RESPONSE pathway of endothelial cells. (**D**) Western blot analysis of total and phosphorylated TBK1 and STING proteins in HPMEC cell lines with or without diABZI treatment (n=3). (**E**) KEGG pathway enrichment analysis of differentially expressed genes in NK cell at 4 h compared to 0 h. (**F**) Transcriptional levels of *Ccl5*, *Cxcl9/10* and *Il15* of HPMEC with or without diABZI or inhibitor treatment (n=3). (**G**) Transcriptional levels of *Ccl4/5*, *Cxcl9/10* and *Il15* of HPMEC with or without diABZI or Compound 53 treatment (n=6). (**H**) Transcriptional levels of *Ccl5* and *Cxcl10* of NCG mice with or without diABZI treatment (n=6). (**I-J**) Flow cytometry counting of NK-92 or hPBMC-NK cells recruited to the basement membrane by HPMECs (n=3). The data are presented as the mean ± SEM. * p < 0.05; ** p < 0.01; *** p < 0.001; ns not significant by ns not significant by unpaired t test or ANOVA followed by Tukey’s multiple comparisons test.

To determine whether endothelial cells could be directly activated by STING agonists, we directly treated human pulmonary microvascular endothelial cell line (HPMEC) with diABZI. Western Blot analysis confirmed that at working concentrations, diABZI significantly increased the phosphorylation levels of STING and TBK1 (Fig. 3D). Validation in mouse cell lines revealed that diABZI directly activated mouse pulmonary endothelial cells (MPMVEC-SV40) and upregulated chemokines (Fig. S4A).

Next, to validate in vitro whether STING activation promotes endothelial cell chemotaxis and inflammatory cytokine secretion, and to investigate the underlying mechanisms, we analyzed pathways enriched in endothelial cell differentially expressed genes at 4 h post-treatment compared to 0 h in scRNA-seq data. Differentially expressed genes enriched in NF-κB, MAPK-CREB-TGF-β, and Wnt-related pathways (Fig. 3E). Targeting these enriched pathways, TBK1-IRF3 pathway and LC3B lipidation, we assessed whether cytokine secretion levels were affected by adding pathway-specific inhibitors to STING activation. Results indicated that the secretion of nearly all factors depended on IRF3 activation. *Cxcl9* expression relied on CREB pathway activation. *Cxcl10* expression was linked to LC3B lipidation and other pathways but independent of NF-κB. *Ccl5* expression mechanisms resembled *Cxcl9*, largely dependent on CREB pathway activation. Whereas *Il15*, similar to *Cxcl10*, was influenced by multiple pathways including LC3B lipidation (Fig. 3F).

Moreover, we were also investigating whether STING’s newly discovered proton pump function was involved in these processes [18]. Compound 53 is a STING agonist that inhibits the function of the STING proton pump [2]. HPMECs activated by diABZI significantly upregulated *Ccl4*, *Ccl5*, *Cxcl9*, *Cxcl10*, and *Il15* expression, whereas Compound 53 administration markedly attenuated the upregulation of *Cxcl10* and *Il15* (Fig. 3G), suggesting that this function depended on the STING proton pump.

In vivo, to confirm this process was indeed independent of other pulmonary immune cells, we modelled NCG immunodeficient mice with equivalent diABZI doses and assessed representative pulmonary chemokine levels. Results showed *Cxcl10* and *Ccl5* remained significantly upregulated (Fig. 3H). To exclude interference from other non-immune cells, we administered identical doses of diABZI to HPMEC, human lung epithelial cells (BEAS-2B), and human lung fibroblasts (HLF-1) in vitro. Endothelial cells exhibited significantly higher cytokine upregulation compared to epithelial and fibroblast cells (Fig. S3B).

### Endothelial cells activated by STING agonists can recruit NK cells

Previously, we observed that within 4 hours of administration, the probability of endothelial cell signal transmission increased substantially, with signals being communicated to NK cells (Fig. 2, F and G). Given that STING agonists promoted endothelial cell expression of chemokines, pulmonary endothelial cells we hypothesized that activated pulmonary endothelial cells may recruit NK cells. Thus, we conducted transwell assays using both the NK-92 cell line and primary NK cells isolated from human PBMCs. Results demonstrated that, compared to the vehicle group, endothelial cells activated by diABZI attracted more NK cells into the lower chamber (Fig. 3, I and J).

### NK cells activated by endothelial cells produce IFN-γ to damage endothelial cells

Previously, we observed that activated endothelial cells could expression NK cell activator IL-15. In subsequent experiments, we investigated the changes occurring in NK cells following signal reception. Therefore, we continued to analyze whether the recruited NK cells would be further activated.

scRNA-seq data revealed that STING expression in NK cells within the lungs rapidly increased within 0–4 hours post-administration, with genes in the IFN-α response pathway significantly upregulated (Fig. 4A) and signal reception intensity substantially elevated (Fig. 2, F and G). To investigate the effects of factors secreted by endothelial cells on NK cells, HPMECs were cultured in medium containing diABZI for 4 hours. NK-92 cells were cultured in this medium for 4 hours, after which they were harvested for gene expression analysis. Results revealed that upon receiving endothelial activation signals, NK-92 cells exhibited markedly elevated expression of *Ccl4*, the classical receptor *Ccr5* for CCL5, the receptor *Il15ra* for IL-15, and *Ifng*. (Fig. 2E) Moreover, direct activation of NK-92 cells with diABZI did not induce this upregulation (Fig. 4B), suggesting that the above changes depended on endothelial cell-secreted factors.

**Fig. 4.**
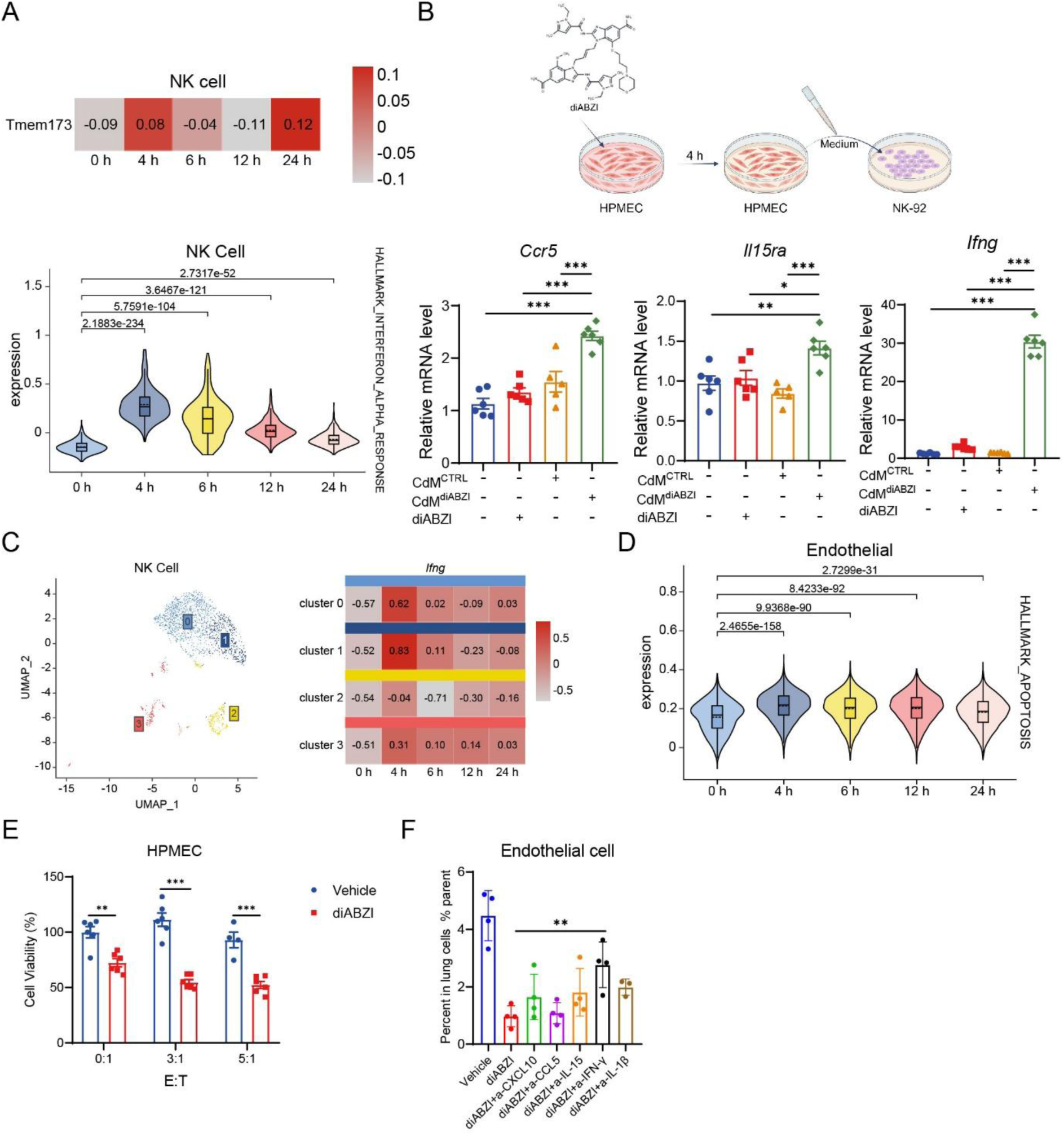
STING-activated endothelial cells release IL-15 to activate NK cells, which produce IFN-γ to damage endothelial cells. (**A**) Heatmap of *Tmem173* expression level and violin plot of gene expression scores for the HALLMARK INTERFERON ALPHA RESPONSE pathway of NK cells at different time points from scRNA-seq data of lung samples. (**B**) Schematic diagram of HPMEC and NK-92 conditioned medium cultivation workflow and transcriptional levels of *Ccr5*, *Il15ra* and *Ifng* of NK-92 (n=6). (**C**) UMAP plots and heatmap of *Ifng* expression level of NK cell subsets for scRNA-seq data of lung samples. (**D**) Violin plot of gene expression scores for the HALLMARK INTERFERON ALPHA RESPONSE pathway of NK cell subsets. (**E**) Relative cell viability after co-culturing NK-92 cells with endothelial cells for 24 hours was determined by measuring the absorbance at 450 nm following incubation with CCK-8 for 2 hours. (**F**) Flow cytometric analysis of endothelial in the lung tissues after neutralization of chemokines and cytokines (n=4). The data are presented as the mean ± SEM. * p < 0.05; ** p < 0.01; *** p < 0.001; ns not significant by ns not significant by unpaired t test or ANOVA followed by Tukey’s multiple comparisons test.

Further analysis of NK cells in scRNA-seq data revealed four distinct subpopulations after dimensionality reduction. Heatmaps indicated that clusters 0 and 1 predominantly comprised *Ifng*-expression populations (Fig. 4C). Literature indicates that IFN-γ promotes endothelial cell apoptosis, compromising the integrity of the endothelial barrier [19, 20]. We hypothesized that activated NK cells could secrete elevated levels of IFN-γ, thereby promoting endothelial cell apoptosis. scRNA-seq data revealed significantly increased expression of endothelial apoptosis pathway genes within 0–4 hours post intraperitoneal administration (Fig. 4D). Following 24-hour co-culture of NK-92 and HPMEC cells at 0:1, 3:1 and 5:1 ratio in vitro, NK-92 cells were removed, and then we add CCK-8 to detect the viability of HPMECs. We observed a significant decrease in relative cell viability of endothelial cells post-co-culture with NK-92 cells, indicating heightened endothelial cell death (Fig. 4E). Then, in vivo, the neutralizing antibodies against CXCL10/CCL5/IL-15/IFN-γ/IL-1β were administered separately with diABZI. Through flow cytometry, results showed that STING agonist reduced the proportion of pulmonary endothelial cells, indicating the impairment of the endothelial barrier. Neutralizing IFN-γ mitigated this reduction, indicating that NK cell activation and the IFN-γ they secrete exert damaging effects on the endothelium (Fig. 4F), demonstrating the important role of IFN-γ, primarily expressed by NK cells, in this process.

Taken together, we discovered that STING agonists could directly activate endothelial cells, promoting their recruitment and activation of NK cells, and then the activated NK cells could induce endothelial cell apoptosis via IFN-γ.

### *Tbx21^+^* NK cells activated by endothelial cells produce chemokines to recruit neutrophils

Further analysis of NK cell alterations following administration of STING agonists, cell interaction data indicated that, the strength of NK cell-to-neutrophil communication progressively increased (Fig. 2G). Concurrently, neutrophil infiltration within lung tissue substantially increased (Fig. 1I and 2A). Previous studies have demonstrated that during acute lung injury, NK cells upregulate T-bet expression and secrete CXCL1/2 to chemotactically recruit CXCR2^+^ neutrophils, thereby exacerbating disease progression [21]. Therefore, we subsequently analyzed the interactions between NK cells and neutrophils.

Analysis of NK cell T-bet (*Tbx21*) expression at different time points post administration, we found that STING agonists significantly upregulated *Tbx21* expression in cluster0/1 NK cells (Fig. 5A). NicheNet analysis of the *Tbx21*^+^ NK population predicted downstream target genes potentially upregulated following NK cell activation by endothelial cell-derived ligands. Circos plot results indicated that cluster0/1 NK cells, upon receiving *Ccl4*, *Ccl5* and *Il15* signals from endothelial cells, leading to increased expression of chemokines *Ccl3*, *Ccl4*, *Cxcl10*, *Ccr5*, and *Ifng* (Fig. 5B). Moreover, within 0-4 hours, differentially expressed genes in cluster 0 NK cells enriched for neutrophil chemotaxis related pathways (Fig. 5C). Thus, single-cell data analysis preliminarily validated our hypothesis.

**Fig. 5.**
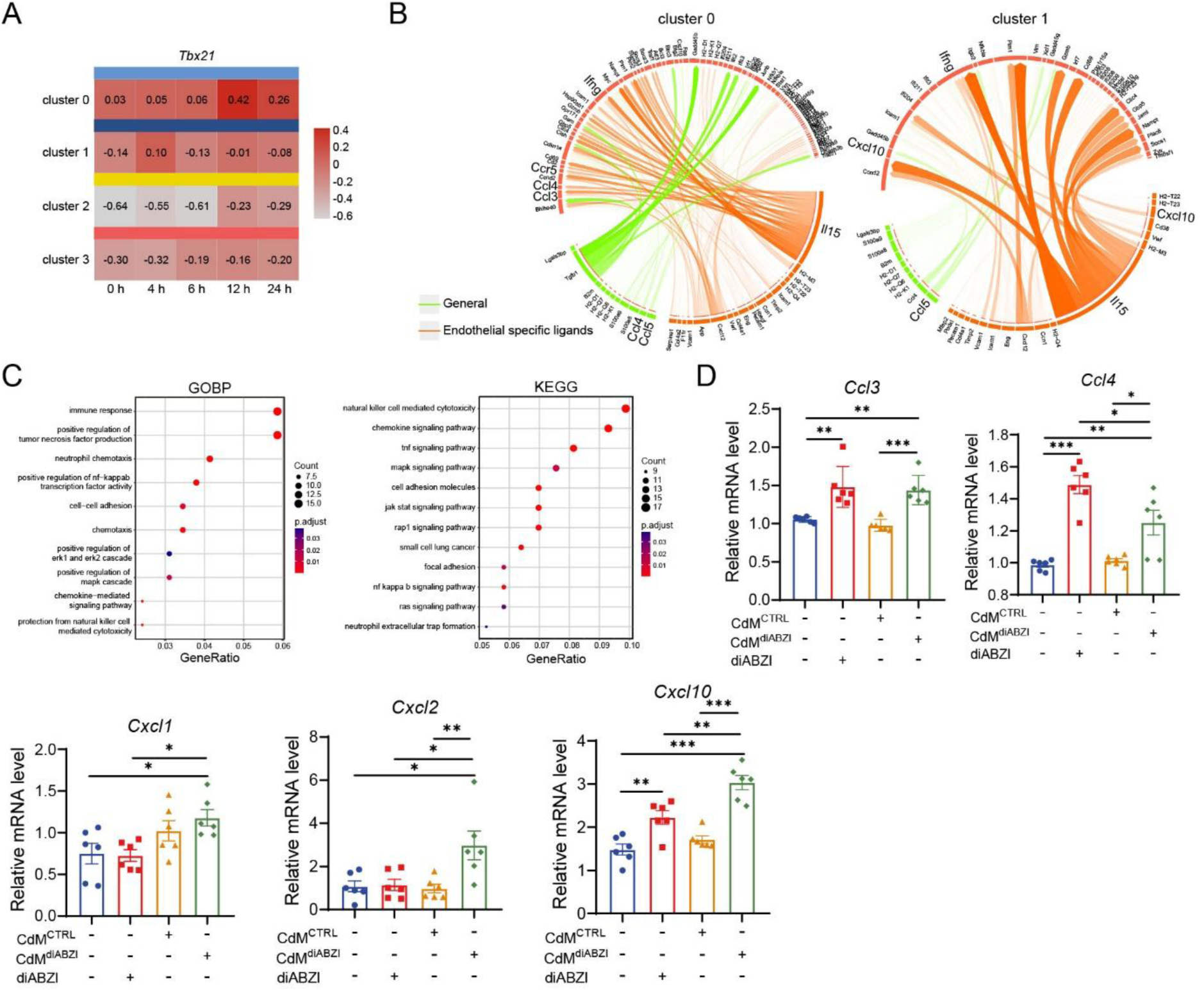
*Tbx21^+^* NK cells activated by endothelial cells produce chemokines to recruit neutrophils. (A) Heatmap of *Tbx21* expression level of NK cell subsets from scRNA-seq data of lung samples. (**B**) Circos plot of NicheNet analysis of the *Tbx21^+^* NK cell population predicted downstream target genes potentially upregulated following NK cell activation by endothelial cell-derived ligands. (**C**) GOBP and KEGG pathway enrichment analysis of differentially expressed genes in cluster0 NK cell at 4 h compared to 0 h. (**D**) Transcriptional levels of *Ccl3/4* and *Cxcl1/2/10* of NK-92 after conditioned medium cultivation with HPMEC (n=6). The data are presented as the mean ± SEM. * p < 0.05; ** p < 0.01; *** p < 0.001; ns not significant by ns not significant by unpaired t test or ANOVA followed by Tukey’s multiple comparisons test.

Subsequently, we assessed the secretion levels of various chemokines by NK-92 cells activated by endothelial cells. Specifically, direct diABZI treatment of NK-92 cells markedly increased *Ccl3/Ccl4/Cxcl10* expression. In contrast, *Cxcl1/Cxcl2/Cxcl10* expression was significantly amplified by endothelial cell activation signals (Fig. 5D).

### Neutrophils secrete IL-1β through positive feedback pathways and form NETs during lung injury

The aforementioned experiments have demonstrated that within a short period following STING agonist administration, NK cells activated by endothelial cells recruit a substantial infiltration of neutrophils into the pulmonary immune microenvironment. Furthermore, neutrophil signaling activity peaks at 12 hours (Fig. 2F and G), with IL-1β secretion reaching its highest level at this time point (Fig. 2D). To investigate how recruited neutrophils mediate lung injury, differential gene analysis and pathway enrichment of 12-hour neutrophils from scRNA-seq data were performed (Fig. 6A). Differentially expressed genes were enriched not only in chemotaxis but also in neutrophil extracellular trap (NET) formation and related pathways (Fig. 6B). Previous studies have demonstrated that in acute lung injury in mice, NETs exacerbate disease progression by mediating parenchymal cell death [22, 23].

**Fig. 6.**
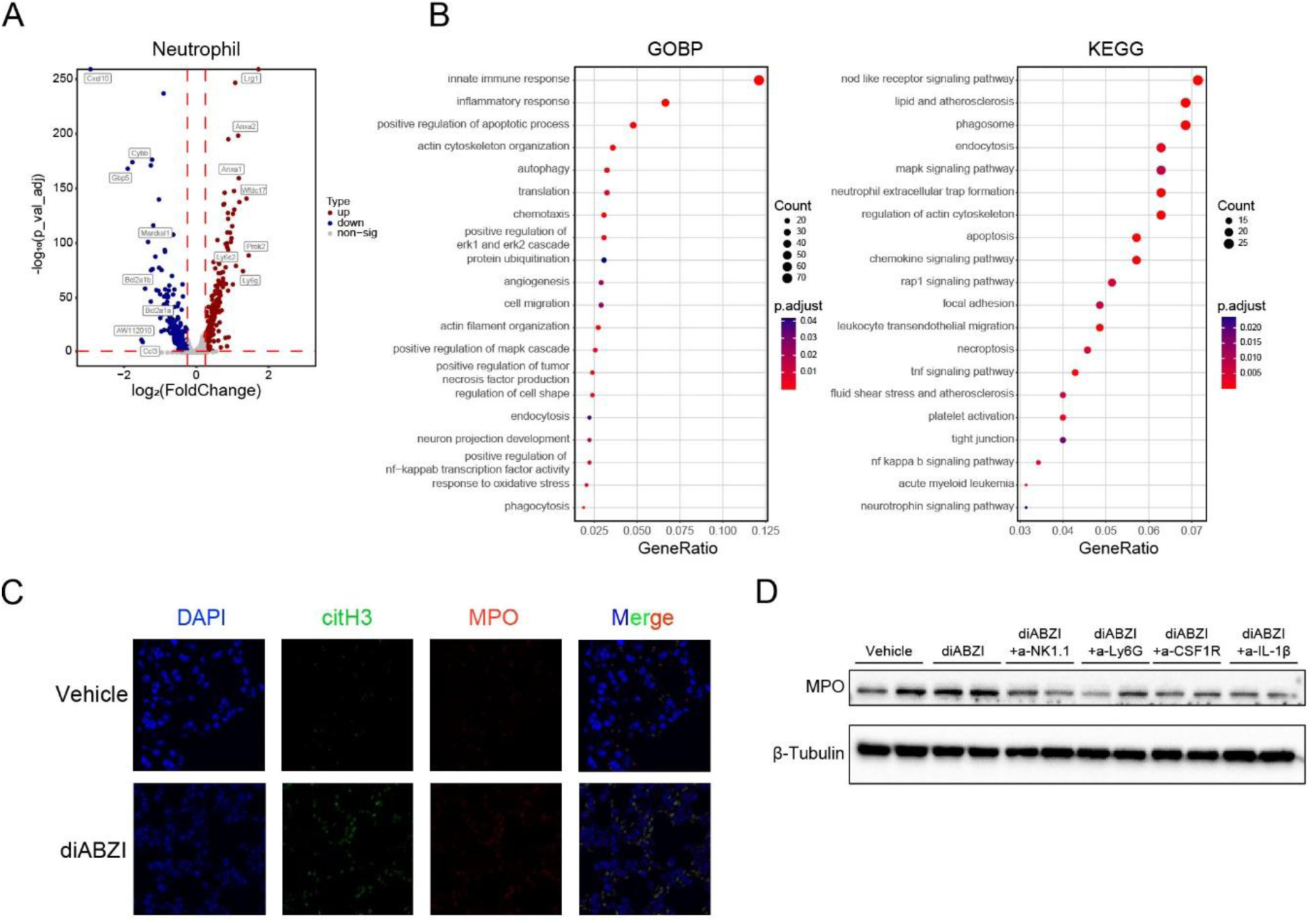
Neutrophils secrete IL-1β through positive feedback pathways and form NETs during lung injury. (**A**) Volcano plot of differentially expressed genes in Neutrophil at 12 h compared to 6 h. (**B**) GOBP and KEGG pathway enrichment analysis of differentially expressed genes in Neutrophil at 12 h compared to 6 h. (**C**) Representative images of IF staining of citH3 and MPO in lung tissues. (**D**) Western blot analysis of MPO in mice lung tissues after depletion of immune cells. The data are presented as the mean ± SEM. * p < 0.05 by unpaired t test or ANOVA followed by Tukey’s multiple comparisons test.

Both multiplex immunofluorescence assays and western blot confirmed increased NETs during STING agonist-mediated pulmonary injury (Fig. 6B and 6C). To validate the contributions of potential immune cells and cytokines, we employed neutralizing antibodies to eliminate NK cells, neutrophils, and interstitial macrophages, and neutralized IL-1β in the diABZI treatment models. Pulmonary tissues expression of the NET marker MPO was assessed by Western blot. Results demonstrated that STING agonists increased pulmonary MPO expression, whilst depletion of NK cells, neutrophils, and interstitial macrophages, and neutralization of IL-1β, all reduced pulmonary MPO levels (Fig. 6C).

Take together, recruited neutrophils may cause lung injury through NETs.

### Immune cells depletion and cytokine neutralization in vivo validates the mechanism of lung injury caused by STING agonist

To validate the contribution of the aforementioned immune cells and cytokines to the STING-mediated toxicity mechanism, we performed immune cell depletion and cytokine neutralization in mice. For neutrophils, which exhibited the most pronounced infiltration increase post-treatment, NK cell depletion reduced their proportion among total lung cells, confirming the chemotactic effect of NK cells on neutrophils during lung injury. Neutralizing IL-1β reduced the increased NK cell infiltration, whereas neutrophil depletion further exacerbated NK cell infiltration. Regarding the markedly reduced pulmonary endothelial cells post treatment, the proportion rebounded following depletion of NK cells, neutrophils, or macrophages, suggesting the role of these three cell types in STING agonist induced pulmonary injury. Neutrophil depletion or IL-1β neutralization partially mitigated the increased interstitial macrophages infiltration, suggesting neutrophils and their secreted IL-1β may exert chemotactic effects on interstitial macrophages (Fig. 7A).

**Fig. 7.**
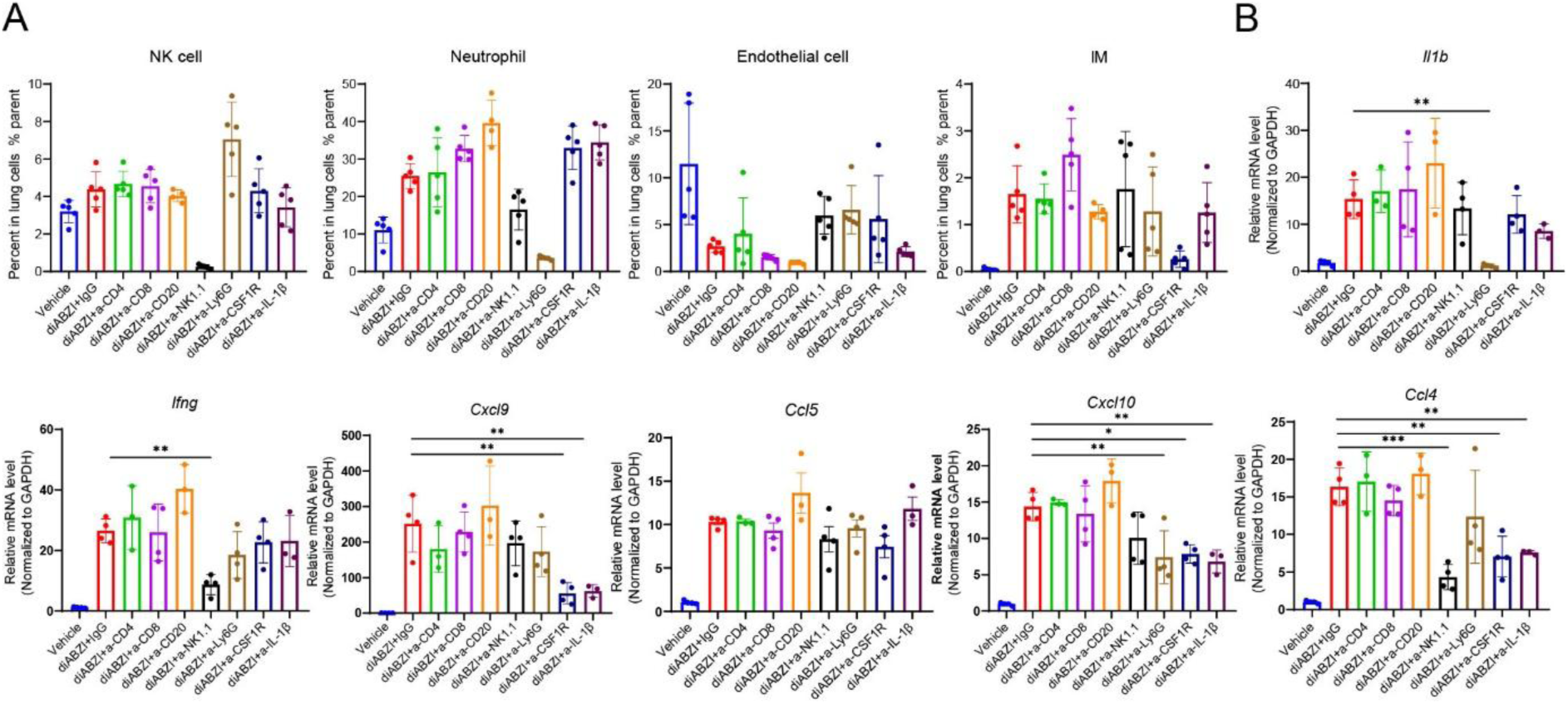
Immune cells depletion and cytokine neutralization in vivo validates the mechanism of lung injury caused by STING agonist. (**A**) Flow cytometric analysis of NK cell, neutrophil, endothelial and IM in the lung tissues (n=4). (**B**) Transcriptional levels of *Il1b*, *Ifng*, *Ccl4/5* and *Cxcl9/10* of the lung tissues after neutralization of chemokines and cytokines (n=5). The data are presented as the mean ± SEM. * p < 0.05; ** p < 0.01; *** p < 0.001; ns not significant by ns not significant by unpaired t test or ANOVA followed by Tukey’s multiple comparisons test.

RT-qPCR analysis of inflammatory cytokines and chemokines in lung tissues confirmed that IL-1β was primarily secreted by neutrophils. Depletion of NK cells and macrophages, alongside IL-1β neutralization, partially alleviated this upregulation. These findings indicated that NK cell depletion indeed reduced neutrophil infiltration, thereby lowering IL-1β secretion levels, with IL-1β secretion potentially exhibiting a positive feedback response.

Regarding IFN-γ, primarily secreted by NK cells, deletion of neutrophils reduced its secretion levels, further validating the interaction between NK cells and neutrophils in the toxic mechanism. Overall, deletion of macrophages, neutrophils, NK cells, or neutralization of IL-1β reduced chemokines production to varying degrees. However, STING agonist-induced *Ccl5* upregulation was largely unaffected by these immune cell deletions, suggesting that *Ccl5* elevation depends on other non-immune cells, including endothelial cells (Fig. 7B).

This directly demonstrated that these immune cells and inflammatory mediators contribute to STING agonist-induced lung injury.

## DISCUSSION

In this study, we have demonstrated the crucial role of the “endothelial-NK-neutrophil” axis in the pulmonary toxicity of STING agonists.

To investigate the effects of STING agonists on the pulmonary immune microenvironment and gene expression, we performed dynamic monitoring of mouse lung tissue at different time points post administration using single-cell RNA-seq and flow cytometry. We observed that STING agonists induced dramatic alterations in the immune microenvironment within a short timeframe. During the early phase, the proportion of T and B lymphocytes plummeted. In fact, this phenomenon has been observed in multiple studies involving gain-of-function STING mutation mice, as well as in experiments involving the administration of STING agonists such as cGAMP [24–26]. Mechanistically, existing studies suggest this is unrelated to interferon signaling or the canonical IRF3 pathway, and may instead be associated with endoplasmic reticulum stress [27–30]. Studies have also indicated that high-dose intratumoral administration of STING agonists can inhibit the expansion of anti-tumor CD8^+^ T cells [31]. This immunotoxicity effect of STING agonists may compromise their sustained antitumor activity.

Cell communication analysis revealed that during the early phase, pulmonary vascular endothelial cells served as the primary signal emitters, with NK cells acting as the core signal recipients. This suggests that endothelial cells, being the first to encounter circulating STING agonists, may be primed for activation and subsequently initiate immune cell recruitment.

The role of endothelial cells in lung injury is gradually being recognized. For example, researchers found that overexpression of cGAS in pulmonary endothelial cells promotes its expression of CCL5 to recruit T cells [32]. These T cells then secrete IFN-γ, causing endothelial damage that mediates the development of pneumonia. Furthermore, multiple studies using STING N153S gain-of-function mice have revealed that sustained STING activation leads to severe pneumonia in these animals [26, 33]. Chimeric mouse experiments indicate that STING activation in endothelial cells initiates pneumonia, but its progression still requires STING activation in immune cells [34, 35].

Both NicheNet predictions from scRNA-seq and in vitro transwell assays confirmed that NK cells activated by endothelial cells exhibit a phenotype characterized by high expression of the transcription factor T-bet (Tbx21), secreting factors such as CXCL1, CXCL2, and CXCL10. These factors strongly recruit neutrophils to lung tissue by acting on receptors including CXCR2 on neutrophil surfaces [21, 36, 37].

Of course, our study has many limitations, and further research is needed. For instances, we have identified the role of NK cells and neutrophils in the pulmonary toxicity of STING agonists, but have not evaluated their function in the antitumor efficacy of STING agonists. This limitation constrains research into the efficacy-toxicity balance of STING agonists. On the other hand, this study did not propose specific STING toxicity intervention strategies. In future studies, we plan to modify STING agonists to avoid their action on endothelial cells, or to combine them with drugs that ameliorate endothelial injury.

In summary, this study systematically characterized the pulmonary toxicity phenotype of STING agonists and elucidated the pathogenesis dependent on the endothelial cell-NK cell-neutrophil axis. Following administration of the STING agonists, pulmonary endothelial cells were initially targeted. Upon activation, they recruited NK cells via CCL5 and further activated NK cells through IL-15. Activated NK cells promoted endothelial cell apoptosis via IFN-γ and recruited neutrophils via the CXCL2-CXCR2 axis. Neutrophils then caused lung injury through NETs, with IL-1β playing a pivotal role in this positive feedback loop. Our research offers insights for developing safer STING agonists in the future, thereby enhancing their potential for clinical application.

## METHODS

### Mice

Male six-to eight-week-old C57BL/6 mice were purchased from Vital River. NCG mice were purchased from GemPharmatech Co., Ltd. All mice were maintained under specific pathogen-free (SPF) conditions in the animal facility of the Shanghai Institute of Materia Medica, Chinese Academy of Sciences. Animal care and experiments were performed in accordance with the Shanghai Institute of Materia Medica, Chinese Academy of Sciences, using protocols approved by the Institutional Laboratory Animal Care and Use Committee (IACUC).

### Establishment of a STING agonist-mediated pulmonary inflammation model

diABZI STING agonist-1 (TargetMol, T11035) was dissolved in DMSO to prepare a 40 mg/mL stock solution. The administration concentration of diABZI is 2 mg/kg (200 μL, dissolved in 40% PEG 300 + 8% Tween-80, then diluted to volume with saline). The pneumonia model was established via intraperitoneal injection on consecutive 3 days (D0-D2). On day D3, blood was collected from the posterior orbital sinus under anesthesia. Following blood collection, mice were euthanized. Lung tissues were rapidly frozen in dry ice for RNA extraction or fixed in formalin solution for paraffin sectioning.

### Cells

NK-92 cell lines were purchased from SUNNCELL and HPMEC cell lines were purchased from IMMOCELL. HPMEC and NK-92 cells were cultured in their respective dedicated media. Human PBMCs were purchased from Milestone. All cells were cultured at 37°C in a 5% CO_2_ humidified atmosphere.

### Immune deletion and cytokine neutralization

In the STING-associated pneumonia model, anti-mouse immune cell deletion antibody (200 μg/animal; Starter) was administered intraperitoneally on days D-1 and D1. On day-1, administer cytokine-neutralizing antibody (200 μg/animal; BioXcell) via intraperitoneal injection. On days 0, 1, and 2, administer cytokine-neutralizing antibody (100 μg/animal; BioXcell) via intraperitoneal injection to maintain neutralizing effects.

### Immunophenotype analysis

After obtaining the whole mouse lung, mince it and place it in digestion buffer (3 mL phenol red-free RPMI 1640 + 2.1 mg collagenase I). Shake at 37°C and 220 rpm for 30 minutes. Subsequently, filter the digested solution through a 70 μm filter membrane. Centrifuge, then lyse with erythrocyte lysis buffer at room temperature for 8 minutes. Filter again after lysis, then centrifuged, washed, and resuspended to obtain a single-cell suspension. Then the cells were blocked with 4% FBS and anti-CD16/32 (BD Biosciences), incubated with surface marker antibodies for 20 minutes at 4℃. Flow cytometry analysis was performed using ACEA NovoCyte and data processing was done through NovoExpress software. Antibody staining was performed following the manufacturer’s recommendations.

### Immunofluorescence

NETs visualization was performed using immunofluorescence confocal microscopy. Formalin-fixed and paraffin-embedded lung specimens from mice were stained with anti-citrullinated histone-3 (citH3, 1:200; Abcam) and anti-myeloperoxidase (diluted 1:200; Abcam 134132), a polyclonal goat anti-mouse Alexa Fluorite 647 antibody (Thermo Fisher) and anti-rabbit Alexa Fluorite 488 antibody (Thermo Fisher) as secondary Abs. The DNA was stained using DAPI (Sigma-Aldrich). All the samples were observed under laser scanning confocal microscopy.

### Cell preparation single cell for RNA-seq

After harvested, lung tissues were washed in ice-cold PBS (Hyclone SH30256.01) and dissociated using SeekMate Tissue Dissociation Reagent Kit A Pro (SeekGene K01801-30) from SeekGene as instructions. DNase Ⅰ (Sigma 9003-98-9) treatment was optional according to the viscosity of the homogenate. Cell count and viability was estimated using SeekMate Tinitan Fluorescence Cell Counter (SeekGene M002C) with AO/PI reagent after removal erythrocytes (Solarbio R1010) and then debris and dead cells removal was decided to be performed or not (Miltenyi 130-109-398/130-090-101). Finally fresh cells were washed twice in the RPMI1640 (Gibco 11875119) and then resuspended at 1×10^6^ cells per ml in RPMI1640 and 2% FBS (Gibco 10100147C).

### Single cell RNA-seq library construction and sequencing

Single-cell RNA-Seq libraries were prepared using SeekOne^®^ DD Single Cell 3’ library preparation kit (SeekGene Catalog No.K00202). Briefly, appropriate number of cells were mixed with reverse transcription reagent and then added to the sample well in SeekOne^®^ chip S3. Subsequently Barcoded Hydrogel Beads (BHBs) and partitioning oil were dispensed into corresponding wells separately in chip S3. After emulsion droplet generation reverse transcription were performed at 42℃for 90 minutes and inactivated at 85℃ for 5 minutes. Next, cDNA was purified from broken droplet and amplified in PCR reaction. The amplified cDNA product was then cleaned, fragmented, end repaired, A-tailed and ligated to sequencing adaptor. Finally, the indexed PCR were performed to amplified the DNA representing 3’ polyA part of expressing genes which also contained Cell Barcode and Unique Molecular Index. The indexed sequencing libraries were cleanup with VAHTS DNA Clean Beads (Vazyme N411-01), analyzed by Qubit (Thermo Fisher Scientific Q33226) and Bio-Fragment Analyzer (Bioptic Qsep400). The libraries were then sequenced on illumina NovaSeq X Plus with PE150 read length.

### Statistical analysis

The in vivo experiments were randomized but the researchers were not blinded to allocation during experiments and results analysis. Statistical analysis was performed using GraphPad Prism 8 Software. A Student’s t test was used for comparison between the two groups. Multiple comparisons were performed using one-way ANOVA followed by Tukey’s multiple comparisons test or two-way ANOVA followed by Tukey’s multiple comparisons test. Detailed statistical methods and sample sizes in the experiments are described in each figure legend. All statistical tests were two-sided and P-values < 0.05 were considered to be significant. ns not significant; *p < 0.05; **p < 0.01; ***p < 0.001.

## Data availability

All data are available in the main text or the supplementary materials. Correspondence and requests for materials should be addressed to Y.L.

## Supporting information

MainText

## Acknowledgments

This work was supported by the Shanghai Rising-Star/Sailing Program (24YF2756000). All the schematics are were created with BioRender.com.

## Author contributions

Conceptualization, L.G. and Y.L.; Methodology, Y.L. and C.C.; Formal Analysis, Y.L.; Investigation, C.C., Y.Z., F.D., R.L., X.Z., S.W., Y.W., F.Q., L.C., R.C., and F.L.; Resources, L.G. and Y.L.; Writing – Original Draft, C.C. and Y.L.; Writing – Review & Editing, L.G. and Y.L.; Supervision, L.G.; Funding Acquisition, L.G. and Y.L.

