## Supplementary material for "STING agonist-mediated endothelial cell activation drives NK cells and neutrophils-dependent pulmonary inflammation": MainText

**This file contains:**

Fig. S1. Lung may be a sensitive target organ for toxicity induced by STING agonists.

Fig. S2. STING agonists upregulate chemokine secretion and enhance the strength of cellular communication.

Fig. S3. Endothelial cells secrete chemokines following STING activation by diABZI.

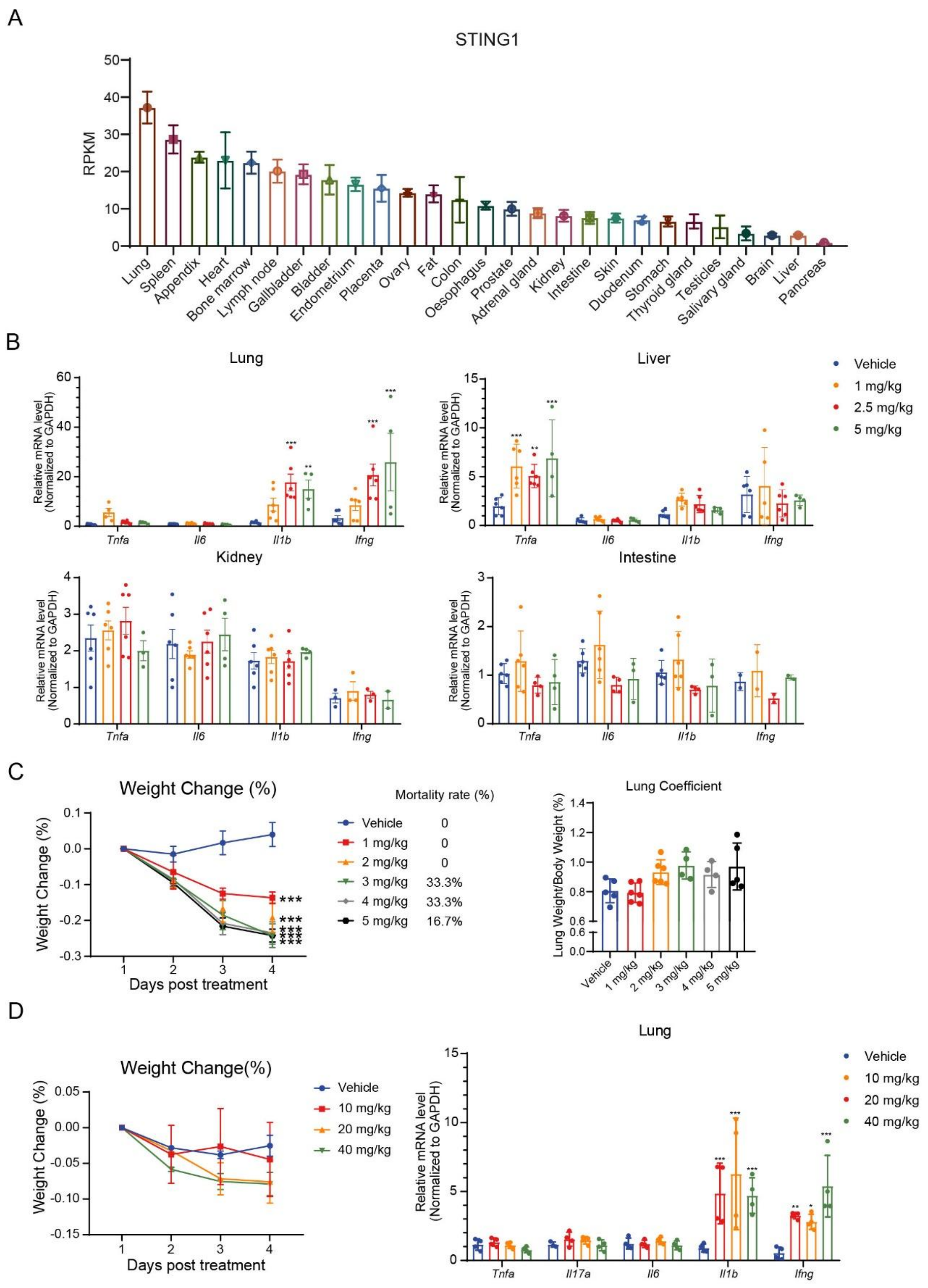


**Fig. S1. Lung may be a sensitive target organ for toxicity induced by STING agonists.** (**A**) Relative expression levels of human STING across various organs. (**B**) Transcription levels of multiple proinflammatory factor in mouse lung, liver kidney and intestine tissues were detected (n=6). (**C**) Body weight change percent of mouse after different doses of diABZI induction from day1 to day4 and lung coefficient on day4 (n=6). (**D**) Body weight change percent of mouse after different doses of 2',3'-

cGAMP treatment from day1 to day4 and transcription levels of multiple proinflammatory factors in mouse lung on day4 (n=4). The data are presented as the mean ± SEM. * $p < 0.05$; ** $p < 0.01$; *** $p < 0.001$ by unpaired t test or ANOVA followed by Tukey’s multiple comparisons test.

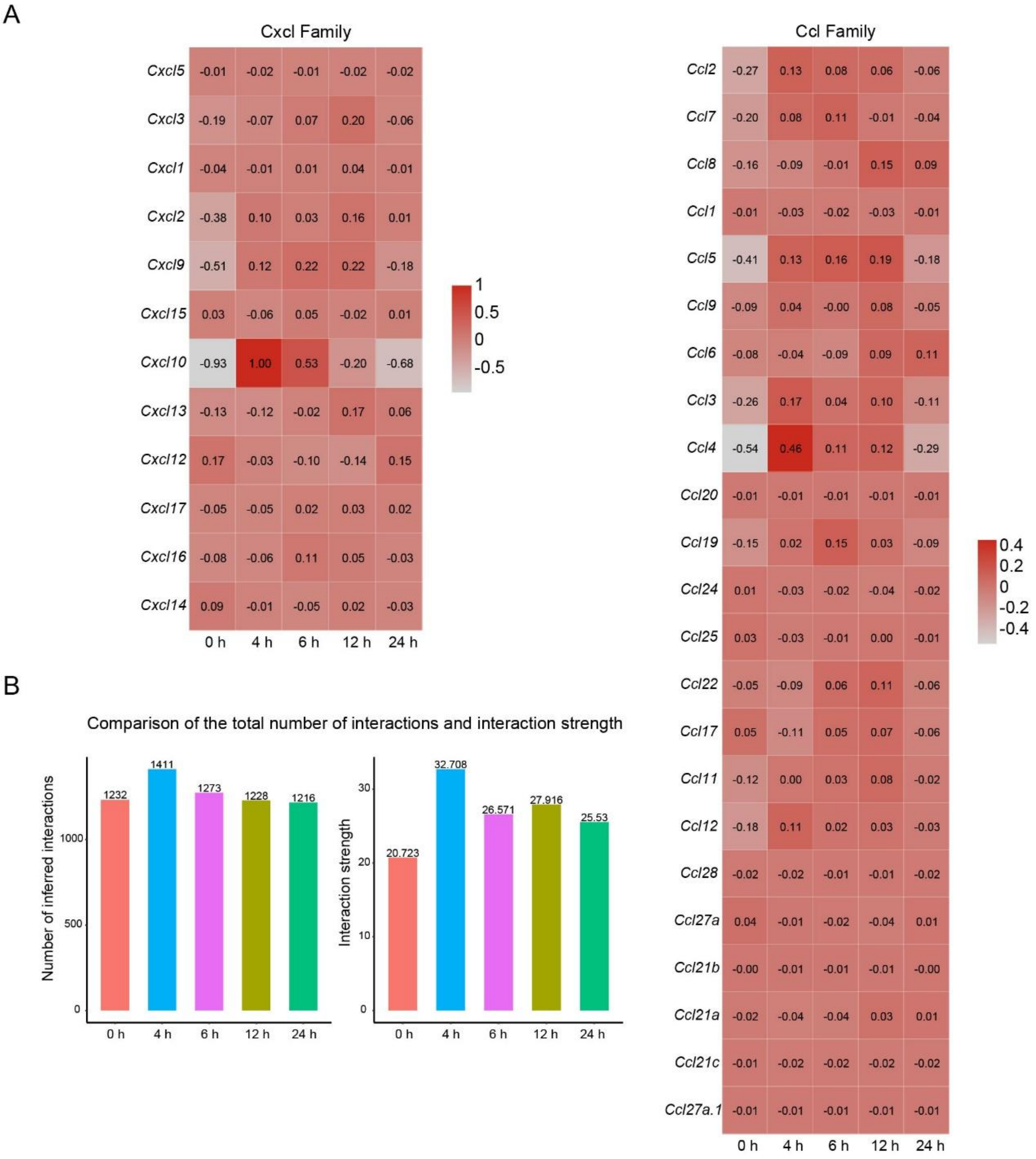


**Fig. S2. STING agonists upregulate chemokine secretion and enhance the strength of cellular communication.** (**A**) Heatmap of chemokine expression levels at different time points from scRNA-seq data of lung samples. (**B**) Intercellular communication strength at different time points from scRNA-seq data of lung samples or ANOVA followed by Tukey's multiple comparisons test.

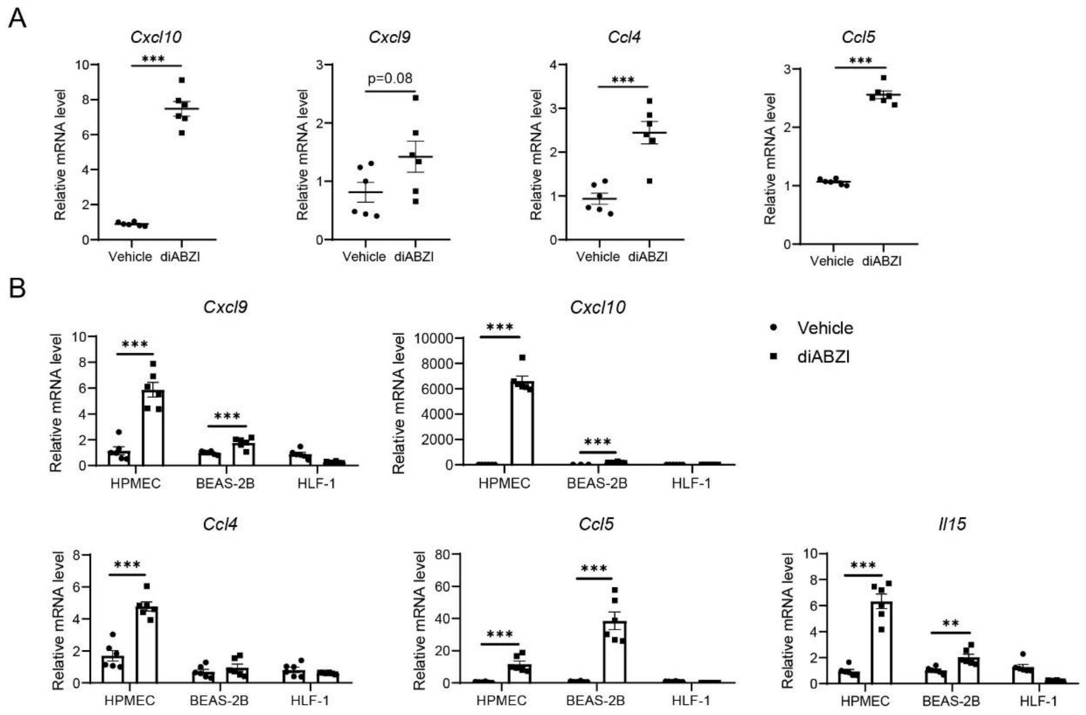


**Fig. S3. Endothelial cells secrete chemokines following STING activation by diABZI.** (**A**) Transcription levels of chemokines of mouse pulmonary microvascular endothelial cells (MPMVEC-SV40) with or without diABZI treatment (n=6). (**B**) Transcription levels of chemokines of HPMEC, normal human pulmonary epithelial cells (BEAS-2B), human lung fibroblasts (HLF-1) with or without diABZI treatment (n=6). The data are presented as the mean ± SEM. ** $p < 0.01$; *** $p < 0.001$ by unpaired t test
